# Constrained inference in sparse coding reproduces contextual effects and predicts laminar neural dynamics

**DOI:** 10.1101/555128

**Authors:** Federica Capparelli, Klaus Pawelzik, Udo Ernst

## Abstract

A central goal in visual neuroscience is to understand computational mechanisms and to identify neural structures responsible for integrating local visual features into global representations. When probed with complex stimuli that extend beyond their classical receptive field, neurons display non-linear behaviours indicative of such integration processes already in early stages of visual processing. Recently some progress has been made in explaining these effects from first principles by sparse coding models with a neurophysiologically realistic inference dynamics. They reproduce some of the complex response characteristics observed in primary visual cortex, but only when the context is located near the classical receptive field, since the connection scheme they propose include interactions only among neurons with overlapping input fields. Longer-range interactions required for addressing the plethora of contextual effects reaching beyond this range do not exist. Hence, a satisfactory explanation of contextual phenomena in terms of realistic interactions and dynamics in visual cortex is still missing. Here we propose an extended generative model for visual scenes that includes spatial dependencies among different features. We derive a neurophysiologically realistic inference scheme under the constraint that neurons have direct access to only local image information. The scheme can be interpreted as a network in primary visual cortex where two neural populations are organized in different layers within orientation hypercolumns that are connected by local, short-range and long-range recurrent interactions. When trained with natural images, the model predicts a connectivity structure linking neurons with similar orientation preferences matching the typical patterns found for long-ranging horizontal axons and feedback projections in visual cortex. Subjected to contextual stimuli typically used in empirical studies our model replicates several hallmark effects of contextual processing and predicts characteristic differences for surround modulation between the two model populations. In summary, our model provides a novel framework for contextual processing in the visual system proposing a well-defined functional role for horizontal axons and feedback projections.

**Author summary:** An influential hypothesis about how the brain processes visual information posits that each given stimulus should be efficiently encoded using only a small number of cells. This idea led to the development of a class of models that provided a functional explanation for various response properties of visual neurons, including the non-linear modulations observed when localized stimuli are placed in a broader spatial context. However, it remains to be clarified through which anatomical structures and neural connectivities a network in the cortex could perform the computations that these models require. In this paper we propose a model for encoding spatially extended visual scenes. Imposing the constraint that neurons in visual cortex have direct access only to small portions of the visual field we derive a simple yet realistic neural population dynamics. Connectivities optimized for natural scenes conform with anatomical findings and the resulting model reproduces a broad set of physiological observations, while exposing the neural mechanisms relevant for spatio-temporal information integration.

## Introduction

Single neurons in the early visual system have direct access to only a small part of a visual scene, which manifests in their ‘classical’ receptive field (cRF) being localized in visual space. Hence for understanding how the brain forms coherent representations of spatially extended components or more complex objects in our environment, one needs to understand how neurons integrate local with contextual information represented in neighboring cells. Such integration processes already become apparent in primary visual cortex, where spatial and temporal context strongly modulate a cell’s response to a visual stimulus inside the cRF. Electrophysiological studies revealed a multitude of signatures of contextual processing, leading to an extensive literature about these phenomena which have been termed ‘non-classical’ receptive fields (ncRFs) (for a review, see Angelucci and Shushruth (2013); Series et al. (2003)). ncRF modulations have a wide spatial range, extending up to a distance of 12 degrees of visual angle Mizobe et al. (2001) and are tuned to specific stimulus parameters such as orientation Sengpiel et al. (1997). Modulations are mostly suppressive Walker et al. (2000), although facilitatory effects are also reported, especially for collinear arrangements where the center-stimulus is presented at low contrast Polat et al. (1998) and for cross-orientation configurations Sillito et al. (1995); Levitt and Lund (1997). However, there is also a considerable variability in the reported effects, even in experiments where similar stimulation paradigms were used: for example, Polat et al. (1998) found iso-orientation facilitation for low center stimulus contrasts, whereas another study Cavanaugh et al. (2002a) did not report facilitation at all, regardless of the contrast level. A further example Sillito et al. (1995) found strong cross-orientation facilitation, while Levitt and Lund (1997) reports only moderate levels of cross-orientation facilitation, if at all. These discrepancies might be rooted in differences between the experimental setups, such as the particular choice of center/surround stimulus sizes, contrasts, and other parameters like the spatial frequency of the gratings, but might also be indicative of different neurons being specialized for different aspects of information integration.

From the observed zoo of different effects in conjunction with their apparent variability, the question arises if explanations based on a unique functional principle could provide a unifying explanation of the full range of these phenomena.

Even though the circuits linking neurons in visual cortex are still a matter of investigation, the nature of their properties suggest the emergence of nCRF phenomena is a consequence of the interplay between different cortical mechanisms Angelucci et al. (2017) that employ orientation-specific interactions between neurons with spatially separate cRFs. Anatomical studies have established that long-range horizontal connections in V1 have a patchy pattern of origin and termination, link preferentially cortical domains of similar functional properties, such as orientation columns, ocular dominance columns and CO compartments Gilbert and Wiesel (1989); Malach et al. (1993); Bosking et al. (1997) and extend up to 8 mm Gilbert and Wiesel (1979, 1989). Although the functional specificity of feedback connections from extrastriate cortex is more controversial, some studies Angelucci et al. (2002); Shmuel et al. (2005) have reported that terminations of V2-V1 feedback projections are also clustered and orientation-specific, providing input from regions that are on average 5 times larger than the cRF. These results make both horizontal and feedback connections well-suited candidates for mediating contextual effects, potentially with different roles for different spatio-temporal integration processes.

Is it possible to interpret the structure of these connections in terms of the purpose they serve?

For building a model of visual information processing from first principles, a crucial observation is that visual scenes are generated by a mixture of elementary causes. Typically, in any given scene, only *few* of these causes are present Simoncelli and Olshausen (2001). Hence, for constructing a neural explanation of natural stimuli, sparseness is likely to be a key requirement. Indeed electrophysiological experiments have demonstrated that stimulation of the nCRF increases sparseness in neural activity and decorrelates population responses, in particular under natural viewing conditions Haider et al. (2010); Vinje and Gallant (2000); Wolfe et al. (2010). Perhaps the most influential work that linked sparseness to a form of neural coding that could be employed by cortical neurons was the paradigm introduced by Olshausen and Field Olshausen and Field (1996). After it was shown that sparseness, combined with unsupervised learning using natural images, was sufficient to develop features which resemble receptive fields of primary visual cortex Olshausen and Field (1996, 1997); Bell and Sejnowski (1997); Rehn and Sommer (2007), a number of extensions have been proposed that have successfully explained many other aspects of visual information processing, such as complex cell properties Hyvärinen and Hoyer (2001) and topographic organization Hyvärinen et al. (2001). Moreover, a form of code based on sparseness has many potential benefits for neural systems, being energy efficient Niven and Laughlin (2008), increasing storage capacity in associative memories Baum et al. (1988); Charles et al. (2014) and making the structure of natural signals explicit and easier to read out at subsequent level of processing Olshausen and Field (2004). Particularly noteworthy is the fact that these statistical models can be reformulated as dynamical systems Rozell et al. (2008), where processing units can be identified with real neurons having a temporal dynamics that can be implemented with various degrees of biophysical plausibility: using local learning rules Zylberberg et al. (2011), spiking neurons Hu et al. (2012); Shapero et al. (2013) and even employing distinct classes of inhibitory neurons King et al. (2013); Zhu and Rozell (2015). In summary, sparse coding models nicely explain fundamental properties of vision such as classical receptive fields.

But can these models also explain signatures of contextual processing, namely non-classical receptive fields?

Recently, Zhu and Rozell reproduced a variety of key effects such as surround suppression, cross-orientation facilitation, and stimulus contrast-dependent ncRF modulations Zhu and Rozell (2013). In their framework, small localized stimuli are best explained by activating the unit with the optimal match between its input field (‘dictionary’ vector). If the stimuli grow larger, other units become also activated and compete for representing a stimulus, thus inducing ncRF modulations. This mechanism is similar to Bayesian models in which contextual effects are caused by surround units ‘explaining away’ the sensory evidence provided to a central unit Lochmann et al. (2012). The necessary interactions between neural units are mediated by couplings whose strengths are anti-proportional to the overlaps of the units’ input fields. However, most of the effects observed in experiments are caused by stimuli extending far beyond the range of the recorded neuron’s input fields Polat et al. (1998); Walker et al. (2000); Mizobe et al. (2001). Hence the mechanism put forward by this model Zhu and Rozell (2013) can only be a valid explanation for a small part of these effects, covering situations in which the surround is small and in close proximity to the cRF. This observation raises the important question, how sparse coding models have to be extended to better reflect cortical dynamics and anatomical structure. In particular, such models would have to allow for direct interactions between non-overlapping input fields.

If these models are then learned from natural images, which local and global coupling structures emerge, how do they compare to anatomical findings, and do they still exhibit the expected cRF properties? Can inference and learning dynamics be implemented in a biophysically realistic manner? Are such models capable of providing satisfactory explanations of ncRF phenomena, and what are the underlying mechanisms? And finally, which predictions emerge from modeling and simulation for experimental studies?

In this paper, we address the above questions by building a novel framework to better capture contextual processing within the sparse coding paradigm. In particular, we define a generative model for visual scenes that takes into account spatial correlations in natural images. To perform inference in this model, we derive a biologically inspired dynamics and a lateral connection scheme that can be mapped onto a neural network of populations of neurons in visual cortex. We show that the emerging connectivity structures have similar properties to the recurrent interactions in cortex. Finally, we evaluate the model’s ability to predict empirical findings reported in a set of electrophysiological experiments and we show that it replicates several hallmark effects of contextual processing. In summary, our model provides a unifying framework for contextual processing in the visual system proposing a well-defined functional role for horizontal axons.

## Results

### Extended generative model

The low-level, pixel representation of a natural image is multidimensional and complex. However, the corresponding scene can often be described by a much smaller number of high-level, spatially extended components such as textures, contours or shapes, which in turn are composed of more elementary, localized features such as oriented lines or grating patches. For constructing an extended generative model of natural scenes (Fig. 1) we thus assume that high-level components located at position ***r*** in a visual scene (e.g. objects ‘donkey’ or ‘wall’ in Fig. 1) imply the presence of a mixture of more elementary features *i* described by coefficients *a*_*i*_(***r***). For capturing correlations between parts of a spatially extended image component, we further assume that the presence of a feature *i* at one particular location ***r*** can be ‘explained’ by the presence of features *j* at other locations ***r’*** via coefficients *c*_*ij*_(***r***, ***r’***) – e.g. an oriented edge that belongs to a contour entails the presence of a co-aligned edge in its proximity Williams and Thornber (2001); Ernst et al. (2012).

**Figure 1.**
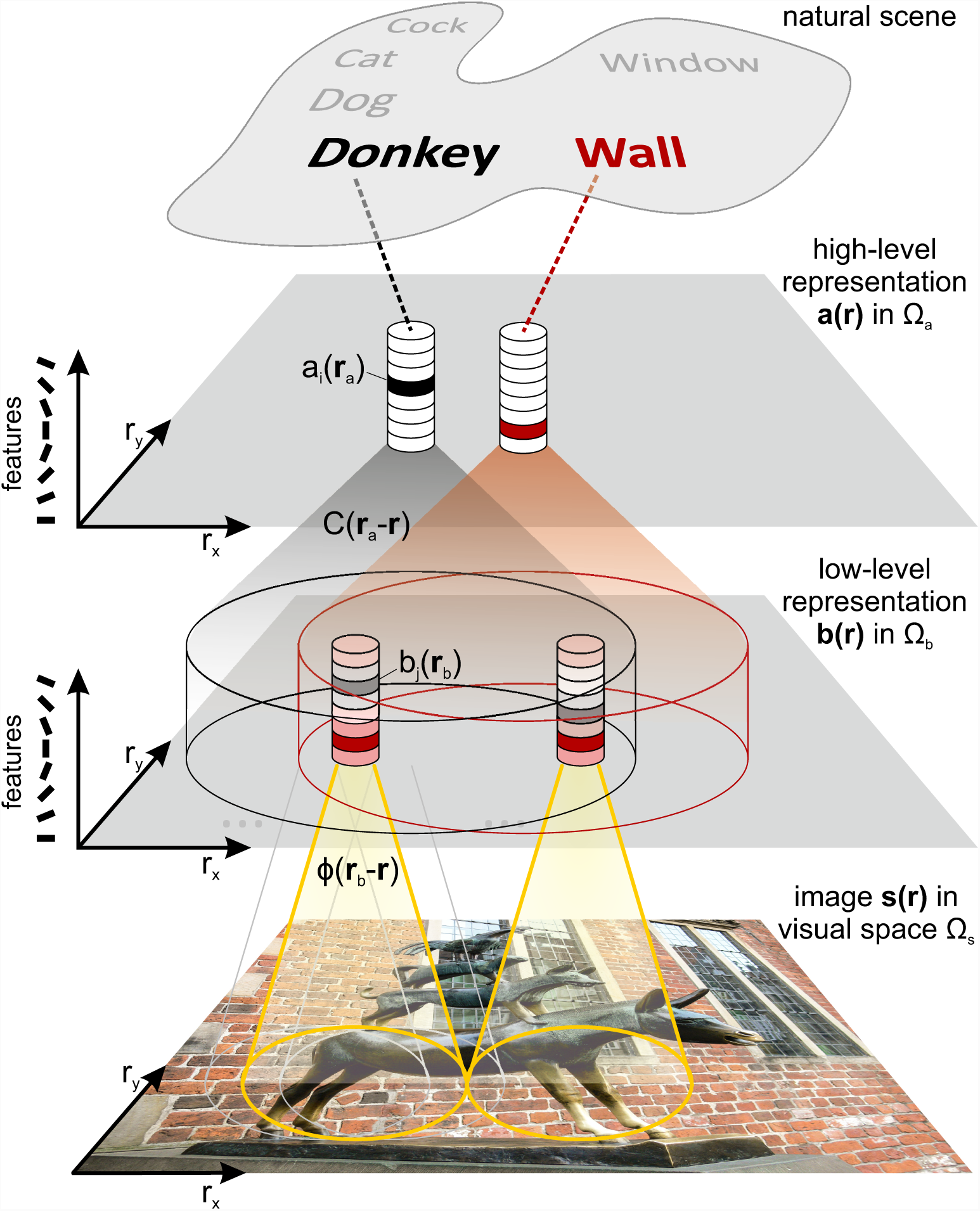
Generative model of natural scenes. The content of a visual scene can be described by the objects that compose it – for instance a window, a wall of a building or the statue of a donkey. Such objects imply the presence of particular shapes, textures and contours, such as the vertical legs of the donkey *a*_*i*_(***r***_*a*_) or the horizontal bricks of the wall. These components extend over large regions in the visual field and hence induce long-range spatial correlations *C*(***r***_*a*_−***r***) between more elementary features (black and red shadings and columns). The feature representation ***b*** thus emerges as a superposition of features of both local and more distant image components. The visual image *s*(***r***) is finally generated from the feature representation by linear superposition of their (pixel) representations or dictionary Φ.

Mathematically, we implement this model by defining

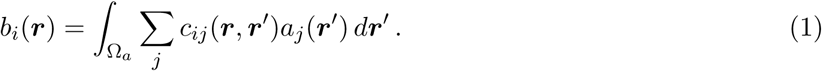

By assuming translation invariance of the *c*_*ij*_ and introducing vector notation we obtain

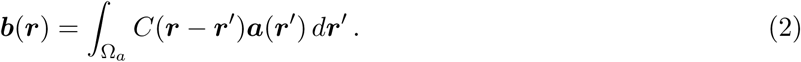

The visual image *s*(***r***), for ***r*** *∈* Ω_*S*_, is finally generated from the feature representation contained in ***b*** by linear superposition

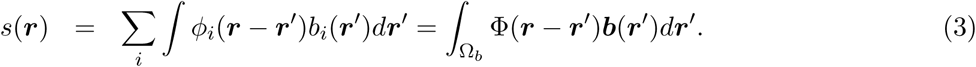

The sets Ω_*a*_, Ω_*b*_ and Ω_*s*_ denote the spatial domains of the representations in ***a*** and ***b***, and the image, respectively. Since we will interpret Ω_*s*_ as the visual field and link ***a*** and ***b*** to cortical representations in the following paragraphs, we will have Ω_*a*_ = Ω_*b*_ = Ω_*s*_ if we assume a 1:1 topographic mapping.

For optimally representing natural scenes, the spatial correlations *C*(***r*** − ***r***^*’*^) and the feature vectors (or dictionary) Φ(***r***) need to be learned from an ensemble of natural images. Since the parameter space for this general spatially extended model is huge, we consider in the following a simplified scenario:

a. Features Φ_*i*_ do not extend over a maximum range *r*_max_ (Fig. 1, indicated by extension of the yellow cones). This is also an implicit assumption in other sparse coding models Olshausen and Field (1996); Lewicki and Sejnowski (2000); Karklin and Lewicki (2003); Rehn and Sommer (2007) which restrict sparse coding to small patches of much larger scenes. Their limited extent is also plausible when features are interpreted as being represented by the synaptic input fields of cortical neurons.
b. *C* is intended to capture long-range spatial correlations extending beyond the range *r*_max_ of elementary features (Fig. 1, indicated by the extension of the black and red cones). The simplest scenario in which *C* plays a role is thus to consider two separate, adjacent image patches ***s***^*u*^ and ***s***^*v*^, whose center locations ***r***^*u*^ and ***r***^*v*^ fulfill |***r***^*v*^ *-* ***r***^*u*^| = *r*_max_. We will interpret these two patches as corresponding to a ‘central’ and a ‘contextual’ stimulus, with their associated feature representations ***a***^*u*^, ***a***^*v*^ and ***b***^*u*^, ***b***^*v*^ (Fig. 2**A**).

**Figure 2.**
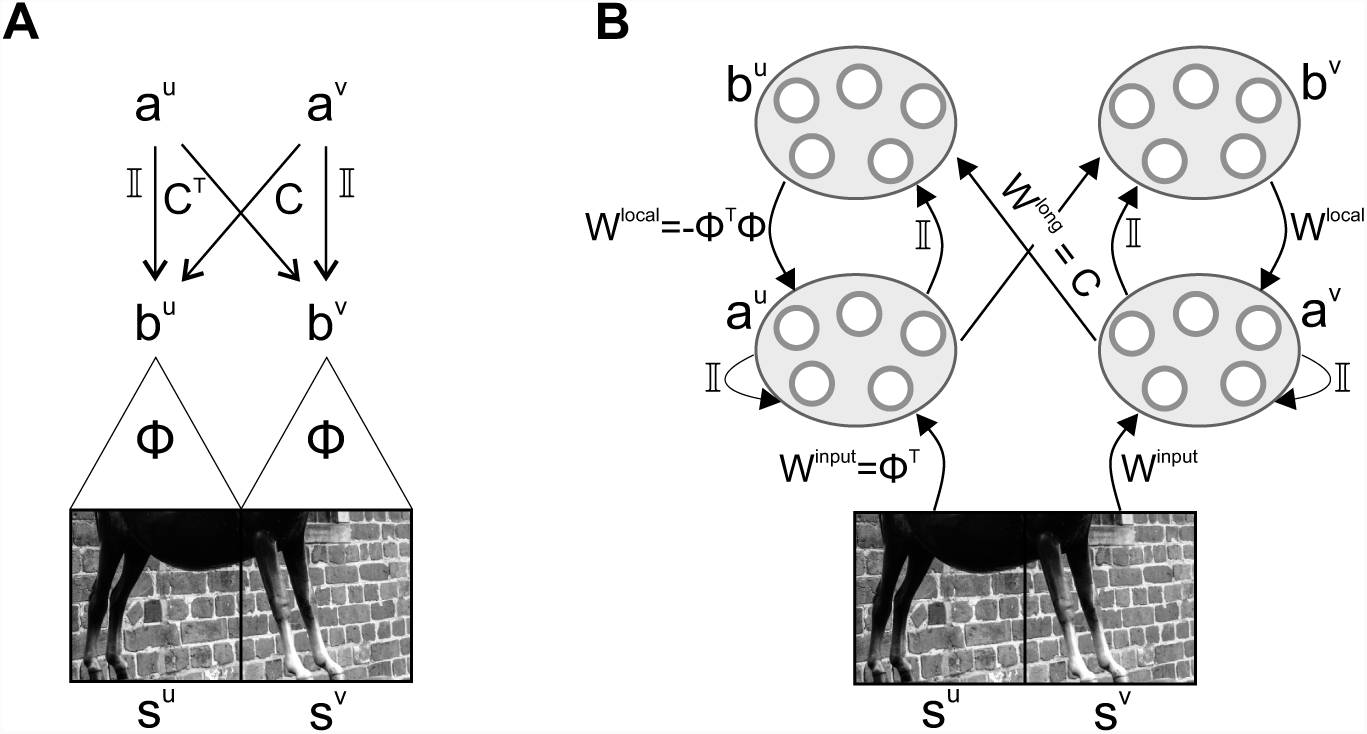
Simplified generative model and neural inference network. (**A**) In a simplified model, we consider visual scenes composed of two horizontally aligned, separate image patches which are encoded by their sparse representation ***a***^*u,v*^ via local features Φ and non-local correlations *C*. (**B**) Inference in the simplified generative model can be performed by a neural population dynamics (21) whose activities represent the coefficients ***a***^*u,v*^ and ***b***^*u,v*^. The corresponding neural circuit involves feedforward, recurrent, and feedback interactions which are functions of the dictionary Φ and of the long-range correlation matrix *C*.

Feature correlations at the same location are set to 1, i.e. *C*^*uu*^ = *C*^*vv*^ = 𝕀. Furthermore, since features separated by a distance *r*_max_ are non-overlapping, the integration in Eq. (3) can be omitted and we arrive at the following set of equations:

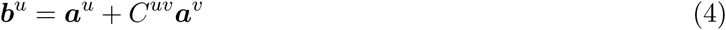

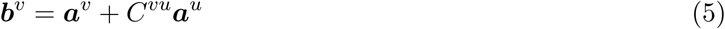

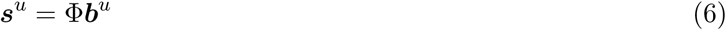

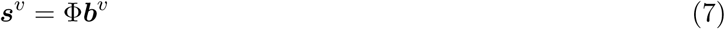

Since we are working with discretized pixel representations of image patches, ***s***^*u,v*^ are now vectors, and Φ becomes a matrix in which the feature vectors are arranged in its columns. Finally, we assume a *reversal symmetry* 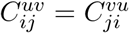 for all *i, j*, which implies *C*^*vu*^ = (*C*^*uv*^)^⊤^: if the presence of a feature *i* at location *u* implies the presence of a feature *j* at location *v*, then the presence of a feature *j* at location *v* should imply the presence of a feature *i* at location *u* to the same extent Williams and Thornber (2001). This allows us to drop the indexes *u, v* and write *C*^*uv*^ = *C* and *C*^*vu*^ = *C*^⊤^. Note that Eq. (6) is identical to standard linear mixture models used to investigate sparse coding of natural scenes Olshausen and Field (1996, 1997); Hoyer (2002). Hence for *C* = 0 different image patches would be encoded independently without using the potential benefits of long-range correlations.

#### Learning visual features and their long-range correlations

To fully define the coding model we posit an objective function, used for optimization of the latent variables and the parameters. In our scheme, it allows to learn which fundamental features Φ_*i*_ are best suited to encode an ensemble of images, and to derive a suitable inference scheme for the latent variables ***a***^*u*^, ***a***^*v*^ such that they optimally explain a given input image (***s***^*u*^, ***s***^*v*^) given the constraints. Most importantly, it allows to determine the spatial relations *C* between pairs of features.

The objective function *E* consists of four terms. The first two quantify how well the two image patches are represented, by means of computing the quadratic error between the patches and their reconstruction. The third and fourth term require the representation in the coefficients ***a***’s and the matrix *C* to be sparse, which is crucial for our assumption that only few non-zero coefficients are necessary to represent a complex image (***s***^*u*^, ***s***^*v*^)_*µ*_ from an ensemble of images *µ* = 1, …, *P*. Mathematically, it is defined by

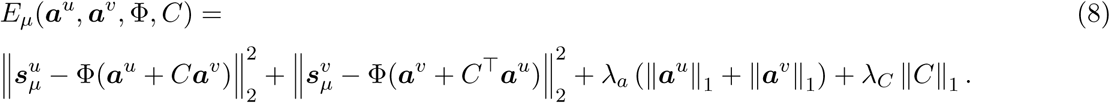

The parameters *λ*_*a*_ and *λ*_*C*_ are sparseness constants, with larger values implying sparser representations. To obtain the matrices Φ and *C* we used a gradient descent with respect to ***a***^*u*^, ***a***^*v*^, Φ and *C* on the objective function defined by equation (8). As image patches ***s***^*u*^, ***s***^*v*^ we used pairs of horizontally aligned, neighboring quadratic patches extracted from natural images (McGill data set Olmos and Kingdom (2004)) after applying a whitening procedure as described in Olshausen and Field (1997). Our optimization scheme consisted of two alternating steps: First, we performed inference for an ensemble of image patches by iterating, for each image *µ*,

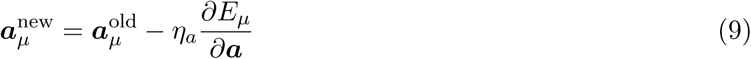

until convergence to a steady state while holding Φ and *C* fixed. Then, we updated Φ or *C* by computing

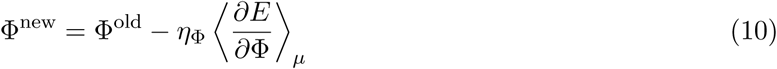

or

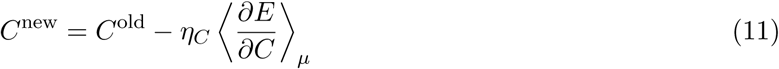

with learning rates *η*_Φ_ and *η*_*C*_, respectively. Angle brackets ⟨· · ·⟩_*µ*_ denotes the average over the image ensemble while keeping the ***a***’s at the steady states (for details, see Methods). This learning schedule reflects the usual assumption that inference and learning take place at different time scales. For increasing computational efficiency, we performed optimization in two phases. First, using only eqs. (9) and (10), we learned the dictionary Φ assuming *C* = 0, and second, using only eqs. (9) and (11), we obtained the long-range correlations *C* while holding Φ fixed.

### Inference with a biologically plausible dynamics

While in theory inference and learning can be realized by the general optimization scheme presented above, in the brain inference needs to respect the neurobiological constraints. In what follows, we derive a dynamics where the mixture coefficients ***a***^*u*^, ***a***^*v*^ and ***b***^*u*^, ***b***^*v*^ are activities of populations of neurons which we hypothesize to realize the necessary computations in cortical hyper-columns connected by local and long-range recurrent interactions (see Fig. 2**B**). Hereby we require populations to have direct access only to ‘local’ image information, conveyed by their synaptic input fields.

For inference, we assume the quantities Φ and *C* to be given and we associate each feature *i* to one neural population having an internal state (e.g. an average membrane potential) and an activation level (i.e., its average firing rate). Following the approach of Rozell et al. Rozell et al. (2008), we define the population activities 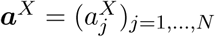 as the thresholded values of the internal states 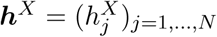 by setting

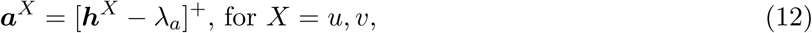

using the sparseness constant *λ*_*a*_ as a threshold, and we let ***h***^*X*^ evolve according to

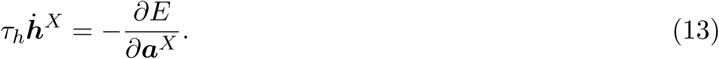

The linear threshold operation ensures the positivity of ***a***, which is a necessary requirement for a neural output. Writing (13) explicitly leads to

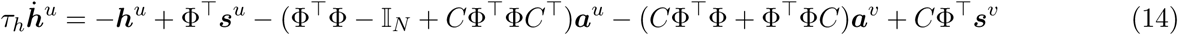

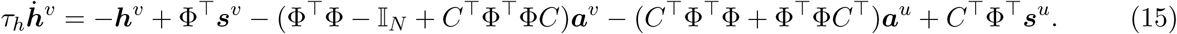

Interpreting these equations in a neural context reveals one problem: The dynamics of the populations at location *u* explicitly depends on the ‘stimulus’ (image patch) at location *v* – and vice versa (last terms on the r.h.s). This dependency violates our assumption of populations having access to only local image information. One way to get rid of this dependence is to approximate the input by its reconstruction suggested by the generative model, that is ***s***^*u*^ = Φ(***a***^*u*^ + *C****a***^*v*^) and ***s***^*v*^ = Φ(***a***^*v*^ + *C*^*T*^ ***a***^*u*^), which leads to

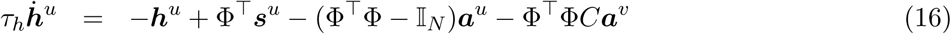

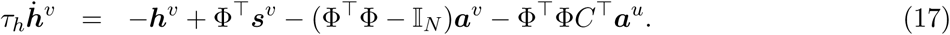

These two equations can be further simplified by extending the dynamical reformulation to include the coefficients ***b*** using eqs. (4) and (5). For this, we define another set of internal variables ***k***^*X*^ satisfying

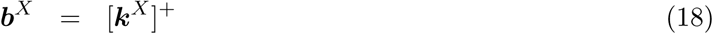

and let them evolve according to a similar relaxation equation (i.e. leaky integration):

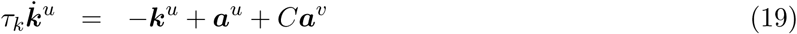

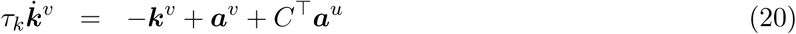

The final model is thus given by the following four differential equations

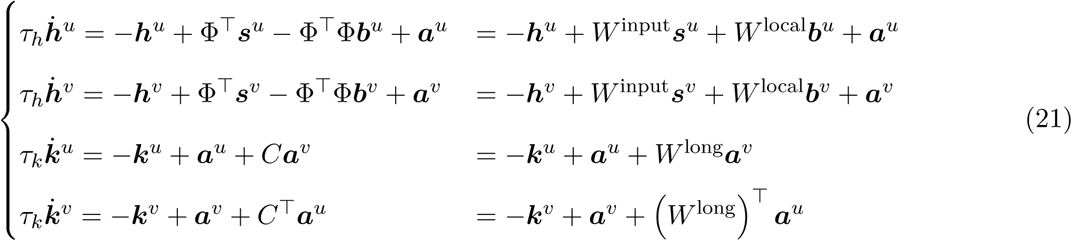

and by the linear threshold operations of eq. (12) and (18).

This temporal dynamics can be implemented in a network of four neural populations organized in two cortical columns (Fig. 2**B**). Specifically, populations ***a***^*u*^ and ***a***^*v*^ in the two columns receive *feed-forward* input *W* ^input^ = Φ^⊤^ from two different locations in the visual field. The input is then processed by a set of recurrent *local* connections that couple population ***a***^*u*^ to ***b***^*u*^ and ***a***^*v*^ to ***b***^*v*^ within the same column (matrices I and *W* ^local^ = −Φ^⊤^Φ). The two populations ***b***^*u*^ and ***b***^*v*^ are also targets of *long-range* connections *W* ^long^ = *C* and (*W* ^long^)^⊤^ originating from populations ***a***^*v*^ and ***a***^*u*^ in the neighboring column, respectively. For example, the two populations ***a*** and ***b*** inside a column could be interpreted as neural ensembles located in different cortical layers, or alternatively as two subpopulations in the same layer, but with different connection topologies. Note that the term ‘long-range’ not necessarily relates to long-ranging horizontal interactions – different anatomical interpretations are possible, and we will speculate on two alternative explanations in the discussion (Fig. 8).

The computation performed within single columns implements a competition based on tuned inhibition between units that code for similar features – which is a typical characteristics of sparse coding models – and it produces a sparse representation of the incoming stimulus. The interactions conveyed by horizontal connections between columns can induce modulatory effects on such a representation. All these connection patterns are completely determined by the matrices Φ and *C*.

Since the representations ***a*** and ***b*** can contain both positive and negative entries, we realized each original unit by two neural populations which we will term ‘ON’ and ‘OFF’ units. Hereby ON-units represent positive activations of the original units, while OFF-units represent negative activations of the original units through positive neural activities. Accordingly, OFF-units are assigned the same shape of the synaptic input field, but with opposite polarity (for implementation details, see Methods). With this necessary extension, equation (13) implies that ***a*** will minimize the energy function *E*: even though the dynamics does not follow the gradient along the direction of its steepest slope, it still performs a gradient descent, since ***a*** is a monotonously increasing function of ***h***.

### Connection patterns and topographies

The link between the formal generative model and its realization as a cortical network allows to interpret Φ and *C* (shown in Fig. 3 and 4) in terms of the connection matrices *W* ^input^, *W* ^local^ and *W* ^long^.

**Figure 3.**
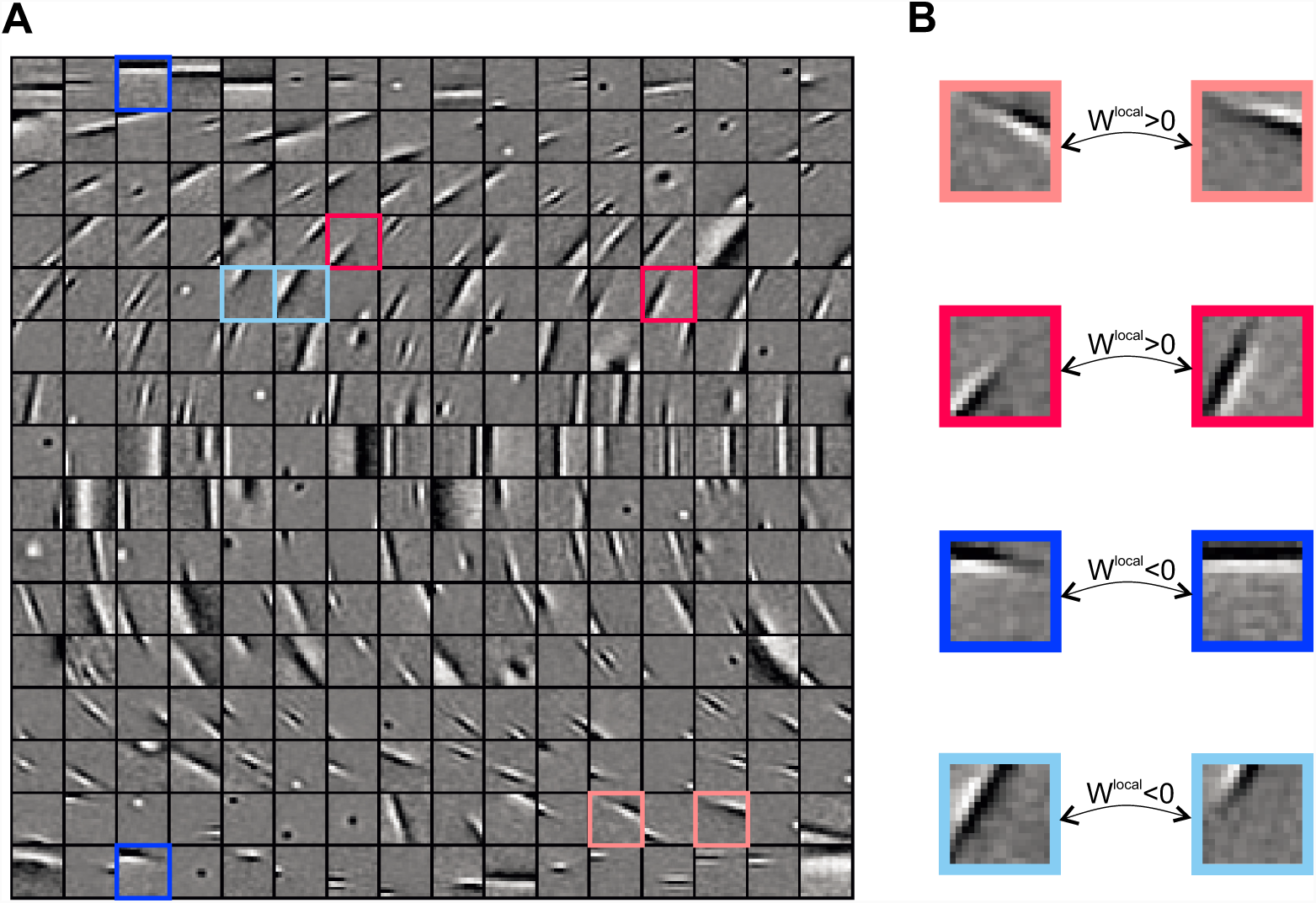
Dictionary Φ and local connections. (**A**) Feature vectors learned by training the model on natural images resemble localized Gabor filters. Features are ordered according to their orientation, which was estimated by fitting a Gabor function. Only a subset of the total set of *N* = 1024 dictionary elements is shown. (**B**) Units with overlapping input fields have strong short-range connections. The sign of the coupling is determined by the arrangement of on/off regions of the input fields: opposite phases correspond to excitatory connections (red) and matching phases to inhibitory connections (blue).

**Figure 4.**
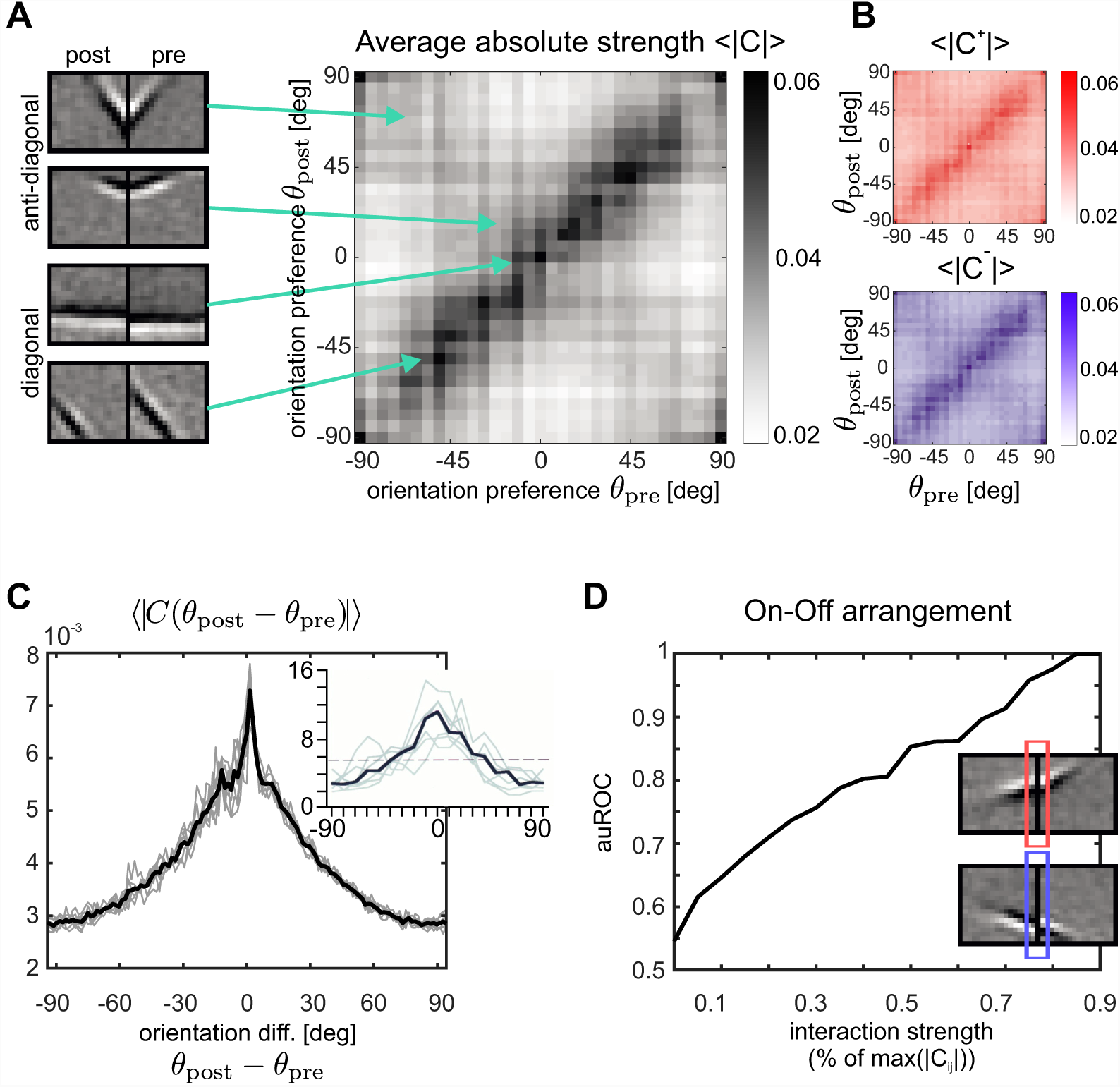
Correlation matrix *C* and long-range interactions. (**A**) Average absolute strength of long-range connections *W* ^long^ = *C* as a function of the orientation preferences of the pre-and postsynaptic units. Each point in the graph represents a connection from a unit responsive to the right portion of the visual field to a unit responsive to the left portion of the visual field (see four examples on the left). (B) Average absolute strength of excitatory (top, red color scale) and inhibitory (bottom, blue color scale) long-range connections as a function of the pre-and postsynaptic orientation preferences as in (**A**). (C) Average absolute strength of long-range connections as a function of the difference in orientation preference of the connected units. For comparison, data from the primary visual cortex of tree shrews are shown in the inset. The graph displays the percentage of boutons contacting postsynaptic sites that differ in orientation preference by a specified amount from the presynaptic injection site of a biocytin tracer. Individual cases are shown in gray and the median is shown in black. The dashed line reflects the percentage of boutons expected in each orientation difference bin if the boutons were distributed evenly over the map of orientation preference (reproduced from Bosking et al. (1997)). (**D**) Long-range interactions between units having positive correlations between the adjacent borders of their synaptic input fields tend to be excitatory (red frame in upper input fields example), while units having negative correlations tend to be inhibitory. This effect increases with increasing absolute coupling strengths |*C*_*ij*_|, as indicated by the area under the ROC curve (auROC) computed from the corresponding correlation distributions for positive and negative connections.

After convergence of the training procedure (see Methods) our model produces feature vectors that resemble Gabor filters (Fig. 3**A**), having spatial properties similar to those of V1 receptive fields. This result is a consequence of the sparseness constraint and does not come as a surprise, since it was obtained in a number of studies before Olshausen and Field (1996); Bell and Sejnowski (1997); Rehn and Sommer (2007) but verifies that our extended framework produces meaningful results by being able to learn similar features. The variety of the dictionary elements is represented in Fig. 3**A** and contains examples of localized and oriented Gabor-like patches, concentric shapes, and structures with multiple, irregularly shaped subfields. Each of the dictionary elements represents the synaptic input field of a neural unit and typically shows up as its classical RF when mapped with localized random stimuli through a reverse correlation procedure Olshausen and Field (1997). For further analysis, we extracted parameters that characterize the cell’s tuning properties – namely its orientation preference, spatial frequency preference, RF center and size – by fitting a Gabor filter to each feature vector (see Methods). Typically, all feature vectors taken together build a complete representation for all orientations (and other stimulus features), thus the columns indicated in Fig. 2**B** are similar to orientation hypercolumns found in primary visual cortex Hubel and Wiesel (1974). The distribution of orientation preferences exhibits a bias for cardinal orientations as observed in physiological studies Wang et al. (2003).

As previously mentioned, short-range interactions are specified by the dictionary matrix through the equation *W* ^local^ = −Φ^⊤^Φ. This implies that the absolute strength with which two units are locally connected is proportional to how closely their respective input fields match. In particular, as it is illustrated in Fig. 3**B**, units with similar orientation preference and opposite phase are excitatorily connected, while units with similar orientation and similar phase are inhibitorily connected.

Together with the dictionary, we also learn the long-range feature correlations *C* (Fig. 4). To investigate which pattern of connections is induced, we computed the average absolute connection strength ⟨|*W* ^long^(*θ*_post_, *θ*_pre_)| ⟩ as a function of the orientation preferences *θ*_post_ and *θ*_pre_ of the units they connect (Fig. 4**A**). The highest absolute connection strengths appear along the diagonal, indicating that pairs of units with similarly oriented input fields tend to be more strongly connected via long-range interactions. The distribution contains another structure, although more faint, located along the anti-diagonal, indicating that pairs of units whose orientations sum up to 0 degrees are also strongly connected. Particular examples of units that have strong long-range coupling are shown in Fig. 4**A**.

This result is consistent with anatomical measurements taken in primary visual cortex of mammals. Several experiments Gilbert and Wiesel (1989); Malach et al. (1993); Weliky et al. (1995); Yoshioka et al. (1996); Bosking et al. (1997) report that horizontal long-range connections in V1 show a ‘patchy’ pattern of origin and termination, linking preferentially cortical domains responding to similar features. We quantified such a tendency in our model by computing the average connection strength as a function of the orientation preference difference Δ*θ* = *θ*_post_ − *θ*_pre_ between pre- and post-synaptic cell. The corresponding graph is shown in Fig. 4**C**, and a similar distribution obtained from anatomical measurements is reported for comparison in the inset.

In three shrew Bosking et al. (1997), cat Schmidt et al. (1997) and monkeys Sincich and Blasdel (2001), it has been shown that long-range connections between neurons of similar orientation selectivity exist primarily for neurons that are retinotopically aligned along the direction of their cells’ preferences. We computed average absolute coupling strength between populations with aligned cRFs (i.e., 0±15 degrees), and between populations with parallel cRFs (i.e., 90 ± 15 degrees), revealing that aligned couplings were indeed 26% percent stronger on average.

When splitting long-range interactions into negative and positive weights, we do not find any significant difference between their dependency on pre- and postsynaptic orientation preference (Fig. 4**B**). However, a different pattern emerges when we take the polarities or phases of the synaptic input fields into account: For this purpose we measured the correlation *ρ* between the right border of the left input field, and the left border of the right input field (colored frames in inset of Fig. 4**D**), which are adjacent in visual space. Excitatory connections tend to exhibit positive correlations, while inhibitory connections tend to exhibit negative correlations. The stronger the couplings, the more pronounced this effect becomes. To quantify this effect, we compared the distributions 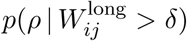 for positive couplings larger than *δ* with the distributions 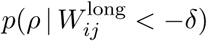 for negative couplings smaller than −*δ* by computing a receiver-operator characteristics ROC). Consistently, we find that separability as quantified by the area under ROC (auROC) increases with *δ* (Fig. 4**D**). This effect is opposite to what we have (by construction) for the short-range connections: while units with similar cRFs *within a column* compete with each other, units with similar cRFs *across two columns* facilitate each other.

### Contextual effects

With the input fields Φ (dictionary) and the long-range interactions *C* obtained from a representative ensemble of natural images, the connectivity of the network represented in Fig. 2 is completely specified. We can then subject the model to arbitrary stimulus configurations and investigate how well the dynamics described by equations (12), (18) and (21) predicts key effects exhibited by real neurons when processing contextual visual stimuli, and whether it can offer a coherent explanation to experimentally established context effects. For this purpose, we first selected units that were well driven and well tuned to the orientation *θ*_*c*_ of small patches *s*_*c*_ of drifting sinusoidal gratings positioned at the center ***r***^*u*^ of the left input region (cf. Fig. 2**B**),

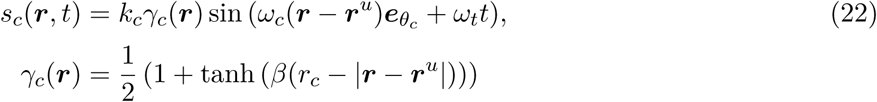

Here *k*_*c*_ denotes grating contrast, *r*_*c*_ the radius of the patch, *ω*_*c*_ its spatial frequency, and *ω*_*t*_ the drifting frequency. *β* controls the steepness of the transition between stimulus und background. Thereby we mimic the situation in experiments in which typically also time-dependent stimuli are used. Subsequently, these selected units were subjected to contextual stimulation, and the induced modulation by the context quantified.

In the following, we will focus on three exemplary stimulation paradigms in contextual processing, assessing size tuning, orientation-contrast effects, and luminance contrast effects.

#### Size tuning

Experiments in monkey and cat Sceniak et al. (1999); Walker et al. (2000) have shown that the stimulation of visual space surrounding the classical receptive field often has a suppressive influence on neurons in V1. Stimuli typically used to reveal this effect consist of a moving grating or an oscillating Gabor patch having the cell’s preferred orientation, and being positioned at the center of its cRF. Recording the neural response while increasing the size of the grating yields the size tuning curve which exhibits two characteristic response patterns Walker et al. (2000), as indicated in Fig. 5**B**: After an initial increase in firing rate with increasing stimulus size, either the cell’s response becomes suppressed and firing rate decreases (upper panel), or firing rate increases further and finally saturates (lower panel). In our model we realized a similar stimulation paradigm by using an optimally oriented grating (eq. (22)) and increasing its size *r*_*c*_. Hence the stimulus first grows towards the border of the input field in which it is centered, and then extends into the neighboring field (i.e., into the right square in Fig. 2**B**). From all selected units, we show the size tuning curves of two exemplary cells in Fig. 5**A**, demonstrating that the model can capture both qualitative behaviors known from cortical neurons.

**Figure 5.**
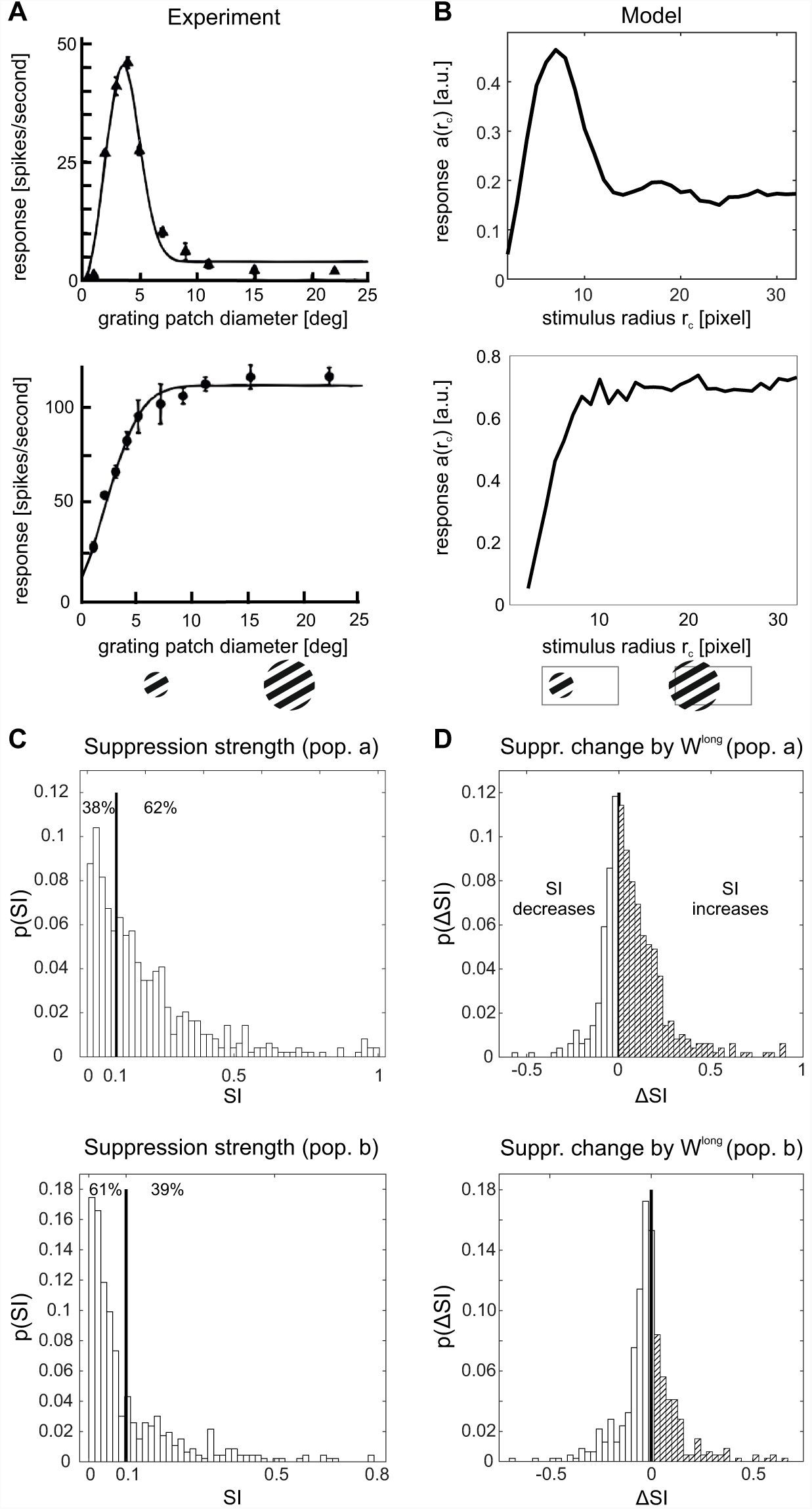
Size tuning and surround suppression. Dependence of neural responses on the size of a circular moving grating presented at the cell’s preferred orientation. (**A**) Single-cell size tuning curves in primary visual cortex of cat exhibiting surround suppression (top) or saturation (bottom). Reproduced from Walker et al. (2000). (**B**) Size tuning curve of exemplary units in the model showing similar behaviour as in (**A**). (**C**) Distribution of suppression indices SI for the full model with long-range interactions. Values of 0 correspond to no suppression, values of 1 to full suppression. (**D**) Change in SI (ΔSI = SI^with long^ − SI^without^) induced by long-range connections. Enhanced suppression occurs more frequently than facilitation in population ***a***, while in population ***b*** one observes the opposite effect.

For quantifying the degree of suppression and the extent to which this effect is present at the population level, we computed for all selected units a suppression index (SI) defined as

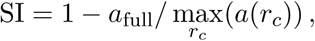

where *a*_full_ was the response to a stimulus fully covering the input field(s). The SI indicates how much, in percentage, the response of a unit at largest stimulus size is reduced with respect to its maximum response, with 0 meaning no suppression and 1 meaning total suppression. The distribution of the SI across all the simulated cells is plotted in Fig. 5**C**. For population ***a***, we find values comparable to what has been found experimentally: Walker et al. (2000) reports that 44% of cells had less than 10% suppression and in the model the percentage of cells with SI*<* 0.1 is 38%. In general, the model shows less suppression (i.e., lower SI values) for population ***b***.

Since surround suppression was already observed in sparse coding models without long-range interactions Zhu and Rozell (2013), we expect this effect to stem from a combination of local and long-range connections. To quantify their roles in producing surround suppression, we simulated a version of the model without long-range interactions by setting *C* = 0. The resulting distribution of changes in SI is shown in Fig. 5**D** and displays a mean increase of the SI for population ***a*** when including long-range connections, indicating that they contribute considerably to suppressive modulation induced by stimuli in the surround. In fact, without long-range interactions the percentage of cells with SI*<* 0.1 becomes 64%, which is quite far from the experimental result reported above. Conversely, the effect of including long-range connections is predominantly facilitatory for population ***b***, leading to a decrease in the observed SI’s.

#### Cross-orientation modulation

Contextual processing is often probed by combining a central grating patch inside the cRF with a surround annular grating outside the cRF. For such configurations, the influence of the surround annulus on the response to an optimally oriented center stimulus was found to be orientation selective. When center and surround have the same orientation, the firing rate modulation is mostly suppressive, as we already know from studying size tuning (previous subsection).

If the surround strongly deviates from the orientation of the center, suppression becomes weaker Levitt and Lund (1997); Sengpiel et al. (1997); Walker et al. (1999); Cavanaugh et al. (2002b) and in some cases even facilitation with respect to stimulation of the center alone is observed Sillito et al. (1995); Jones et al. (2002). In particular, one study in cats Sengpiel et al. (1997) reports three typical response patterns: (I) equal suppression regardless of the orientation of the surround, (II) suppression which decays with increasing difference between the orientations of center and surround, and (III) suppression that is strongest for small differences between orientations of center and surround, and weaker for large orientation differences and orientation differences close to zero. In the literature, the last effect is also termed ‘iso-orientation release from suppression’ (see Fig. 6**A** for examples).

**Figure 6.**
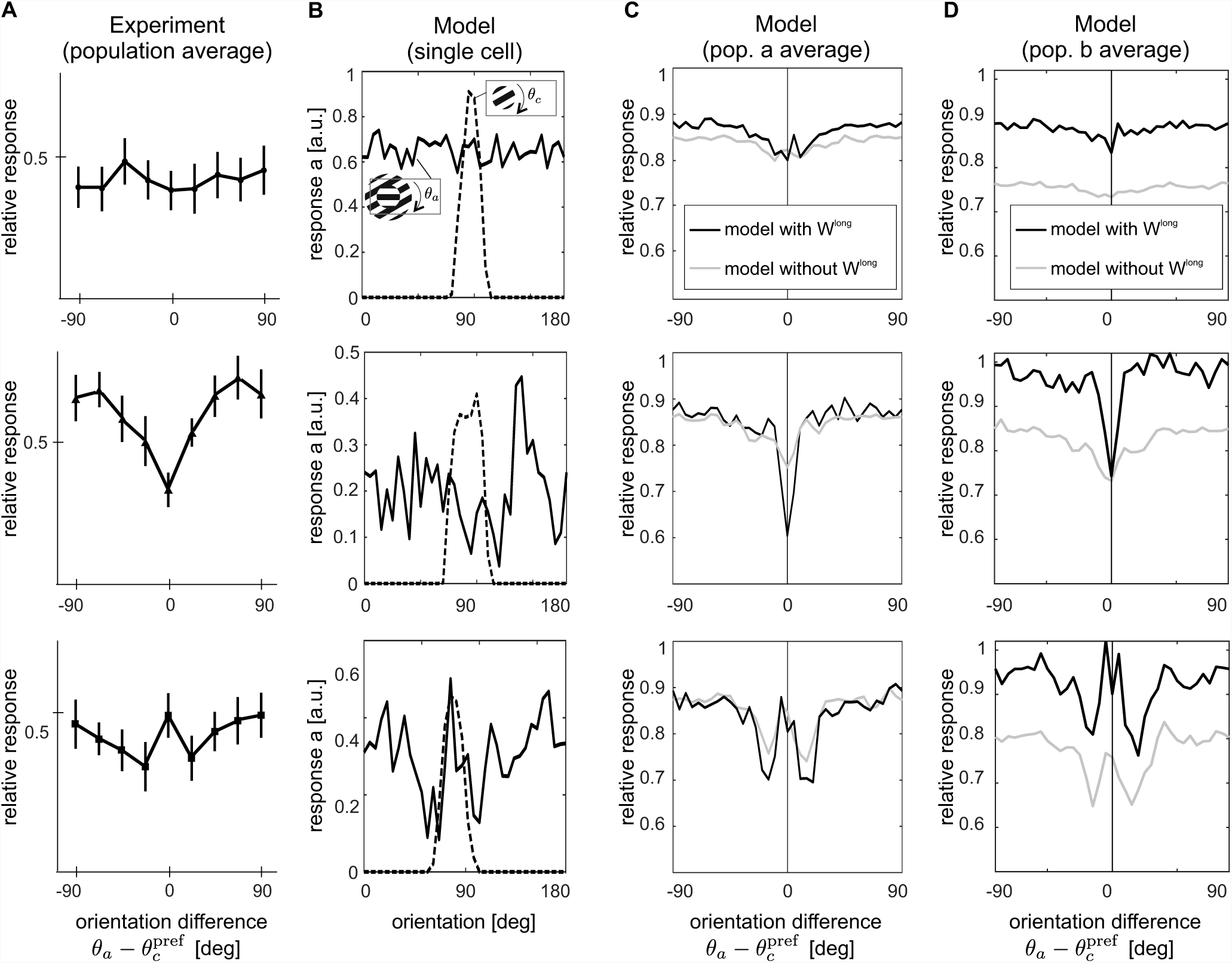
Orientation-contrast modulations. A center stimulus with preferred orientation is combined with an annulus of varying orientations (see insets column (**B**). (**A**) In experiments three response patterns are observed, namely, from top to bottom, untuned suppression, iso-orientation suppression and iso-orientation release from suppression (data replotted from Sengpiel et al. (1997)). The model reproduces these three response patterns both at the single cell level (**B**) and at the population level for ***a*** (**C**) and ***b*** (**D**). For comparison, orientation tuning for a center-alone stimulus is shown by the dashed line in (**B**). In (**C, D**), the gray lines display orientation-contrast tuning of the same ensembles without long-range interactions. Note that in (**A**) and (**C, D**), responses are shown normalized by the response to the center alone at the preferred orientations of the units.

We realized this experimental paradigm in our model by combining a central grating patch (eq. 22) with a surround annulus

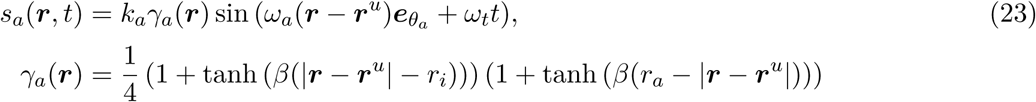

having orientation *θ*_*a*_, spatial frequency *ω*_*a*_ = *ω*_*c*_, inner radius *r*_*i*_ = *r*_*c*_, outer radius *r*_*a*_, and grating contrast *k*_*a*_ = *k*_*c*_. For each neural unit we investigated, the center stimulus had an optimal size defined by the radius *r*_*c*_ for which we obtained the maximum response in the unit’s size tuning curve. The surround annulus had the same parameters as the center patch and extended from the radius of the center patch to the whole input space (as displayed in Fig. 6, stimulus icons in the legends). While the center orientation was held at the unit’s preferred orientation, the surround orientation *θ*_*a*_ was systematically varied between 0 and *π*. For this experiment, we selected all units for which their optimal size was not larger than 21 pixels, to ensure that there was still space for a surround annulus in the restricted input space.

The three distinct behaviors observed in the experiments are qualitatively captured by the model: in Fig. 6**B** (dashed lines) we show the orientation tuning curve of selected units of the model. Adding an annular surround stimulus to an optimally oriented center induces modulations which are mostly suppressive and tuned to the orientation of the surround (Fig. 6**B**, solid lines). Cross-orientation modulations are summarized across the investigated model subpopulation in Fig. 6**C**, **D**, where responses of cells exhibiting the same qualitative behavior are averaged together, as in the experiment (cf. panel **A**, see Methods for a detailed description of the pooling procedure). We distinguish, from top to bottom, untuned suppression, iso-orientation suppression, and iso-orientation release from suppression.

To assess the contributions of long-range connections to these effects, we repeated the experiment with *C* = 0. The population averages over the same categories of behaviors are overlaid in Fig. 6**C,D** in gray. A comparison between the results of the model with and without *C* shows that long-range interactions induce two different effects: enhancing responses for large orientation differences for cells with untuned surround suppression, and increasing maximum suppression for cells with tuned surround suppression in population ***a***. In particular, we observe strong facilitatory effects in population ***b***. This difference between the two populations might explain an apparent contradiction in experimental data where in a similar orientation contrast tuning paradigm one study exhibited strong facilitation Sillito et al. (1995), while a different investigation found only moderate release from suppression Levitt and Lund (1997).

#### Luminance-contrast effects

In addition to orientation, also the relative contrast between the brightness of the center and the surround can be varied. In particular, such stimuli often reveal facilitatory effects, which are more frequently observed when the cRF is weakly activated, for example by presentation of a low-contrast visual stimulus. For many cells in V1 (*≈* 30%, in Polat et al. (1998); Chen et al. (2001)), collinear configurations of center-surround stimuli induce both facilitation and suppression. Here the visual contrast of the center stimulus in comparison to a fixed-contrast surround controls the sign of the modulation, and the point of crossover between suppression and facilitation is related to the cell’s contrast threshold Levitt and Lund (1997); Mizobe et al. (2001); Polat et al. (1998); Sengpiel et al. (1997); Toth et al. (1996). The characteristics of differential modulation is exemplified in Fig. 7**A** where the contrast response function of a single cell in cat V1 (filled circles) is plotted together with the response of the same cell to the compound stimulus (empty circles). The graph shows that the same surround stimulus can enhance the response to a low-contrast center stimulus and reduce the response to a high-contrast center stimulus.

**Figure 7.**
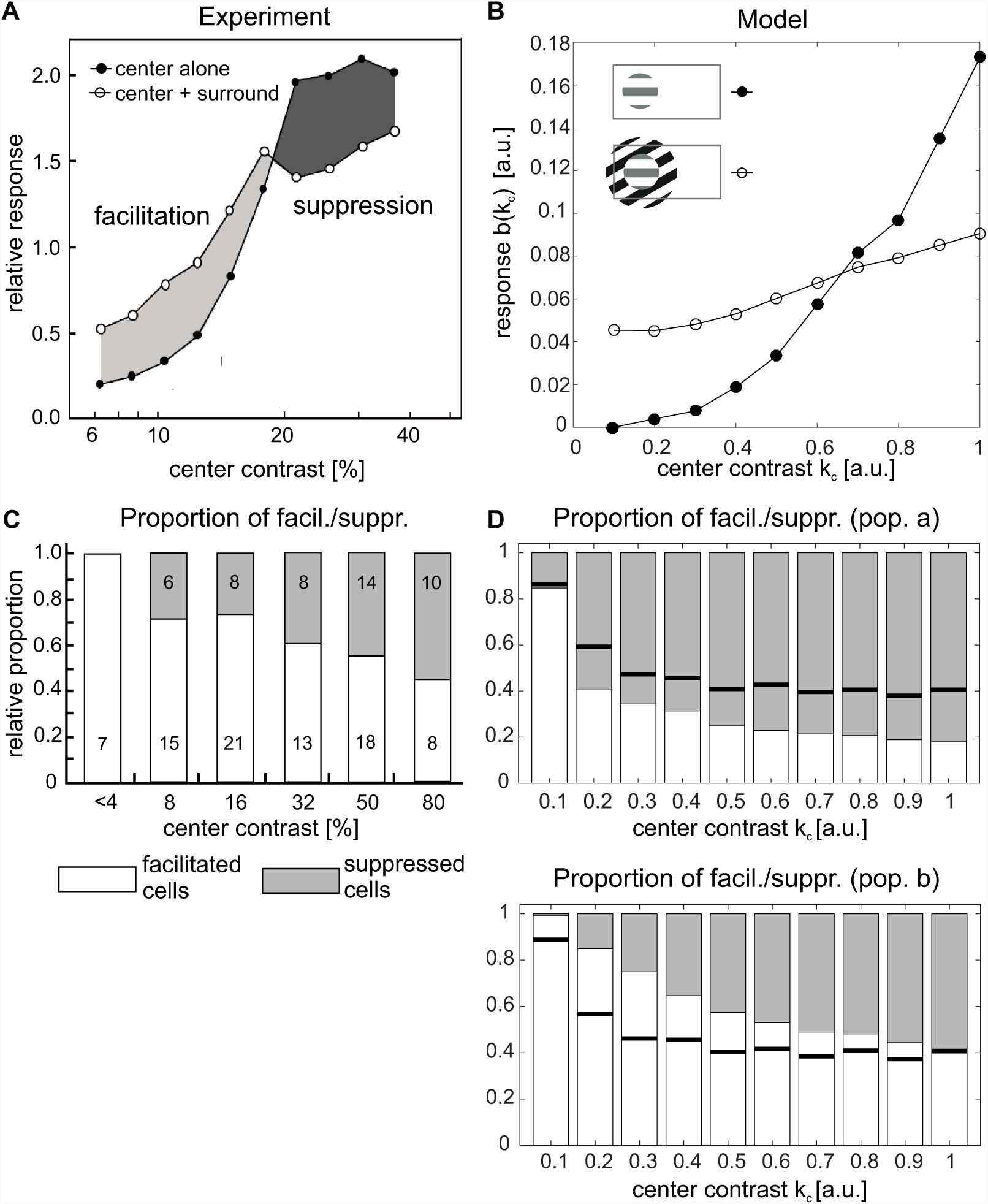
Luminance contrast tuning. **(A, B)** Single-cell responses to a center stimulus of varying contrast without flanking surround stimuli (filled circles) are compared to responses to the same center stimulus combined with high-contrast flanking surround stimuli of the same preferred orientation (open circles) in experiment **(A)** (reproduced with permission from Polat et al. (1998)) and model **(B)**. The stimulus configurations are indicated inside the graphs. **(C, D)** Population statistics, detailing the proportion of cells showing facilitation (light bars) or suppression (gray bars) in dependence on center stimulus contrast. Experimental data in **(C)** is reproduced from Polat et al. (1998), numbers inside the bars indicate the total number of cells showing suppression/facilitation. In the model **(D)**, cells were judged to be significantly facilitated (suppressed) if their activation ratio between center-surround and center alone stimulation *b*^sur^(*k*_*c*_)*/b*^cen^(*k*_*c*_) at contrast *k*_*c*_ was larger than 1 + *ε* (smaller than 1*−ε*), with *ε* = 0.01. Solid black lines indicate proportion of cells showing facilitation without long-range interactions. The top plot in **(D)** shows the statistics for population ***a*** and the bottom plot for populations ***b***.

For obtaining corresponding contrast response curves in our model, we presented each selected unit with a center stimulus of optimal orientation and size of which we varied its contrast *k*_*c*_ (eq. (22)). To mimic the collinear configuration of the compound stimulus, we then placed a surround annulus (eq. (23)) at high contrast *k*_*a*_ = 1, iso-oriented with the center patch (see stimulus icons in Fig. 7), and again varied the contrast of the center patch. The resulting switch from facilitation to suppression, apparent by the crossing of the two response curves, is well captured by the model and illustrated for an example unit in population ***b*** in Fig. 7**B**.

As in previous examples, differential modulation shows considerable variability across recorded cells. In particular, there are V1 neurons which exclusively show suppressive effects, while other neurons exclusively exhibit facilitatory effects. The corresponding statistics is displayed in Fig. 7**C**: For each value of contrast that was tested in Polat et al. (1998), the bars show the proportion of cells that exhibit either facilitation or suppression. In particular, suppression becomes increasingly more common as the contrast of the center stimulus increases. The same analysis applied to our model reveals an identical result (Fig. 7**D**), thus indicating that the model also captures the diversity of behaviors observed in electrophysiology. For population ***b***, the model statistics matches experimental findings also quantitatively. In particular, we observed that the increase in numbers of suppressed cells with increasing center contrast is mainly caused by the long-range connections, since this effect largely disappeared when we set *C* = 0 (horizontal lines in Fig. 7**D**).

## Discussion

The pioneering work of Olshausen and Field Olshausen and Field (1996) demonstrated that simple cell responses in primary visual cortex can be understood from the functional requirement that natural images should be represented efficiently by optimally coding an image with sparse activities. Since then, there have been many attempts to derive also other neuronal response properties in visual cortex from first principles. Common to these models is the framework of generative models, where the activities in an area are considered to represent the results of inference in the spirit of Helmholtz von Helmholtz (1962); Doya et al. (2007). Most of these investigations concentrate on local receptive field properties Olshausen and Field (1997); Bell and Sejnowski (1997); Rehn and Sommer (2007); Hyvärinen and Hoyer (2001); Hyvärinen et al. (2001). More recently, formal models were introduced that can qualitatively reproduce also several established non-classical receptive field effects Karklin and Lewicki (2009); Coen-Cagli et al. (2012); Lochmann et al. (2012); Coen-Cagli et al. (2015) and/or predict interactions resembling features of long-ranging horizontal and feedback connections in cortex Garrigues and Olshausen (2008); Coen-Cagli et al. (2012).

It is, however, unclear how the networks in cortex might perform the inference these models hypothesize given the neurobiological constraints on anatomy and neuronal dynamics. In this regard, the neural implementation proposed by Rozell et al. (2008) provided a significant advance, since it can explain a range of contextual effects Zhu and Rozell (2013) with a neural population dynamics that requires only synaptic summation and can also be extended to obey Dale’s law Zhu and Rozell (2015). But this model still presents a fundamental, conceptual difference to visual cortex: there are no interactions between neurons with non-overlapping input fields and thus the model can not account for the long-range modulatory influences from far outside the classical receptive field.

Here we propose a generative model for sparse coding of spatially extended visual scenes that includes long-range correlations between local patches in natural images. An essential ingredient is the inclusion of plausible neural constraints by limiting the spatial extent of elementary visual features, thus mirroring the anatomical restrictions of neural input fields in primary visual cortex.

### Connection structures

By optimizing model parameters via gradient descent it is possible to determine all connections in the network e.g. from the statistics of natural images. Synaptic input fields Φ resemble classical receptive fields of V1 neurons (Fig. 3**A**). The structure of *C* turns out to have a similar characteristics as the anatomy of recurrent connections in visual cortex, exhibiting a preference to link neurons with similar orientation preferences via long-ranging horizontal axons Kaschube (2014); Schmidt et al. (1997); Gilbert and Wiesel (1989) or via patchy feedback projections Angelucci et al. (2002); Shmuel et al. (2005). Furthermore, we find a bias for collinear configurations being more strongly connected than parallel configurations, matching the observed elongation of cortical connection patterns along the axis of collinear configurations in the visual field in three shrew Bosking et al. (1997), cat Schmidt et al. (1997) and monkeys Sincich and Blasdel (2001). These connection properties reflect regularities of the visual environment such as the edge co-occurrence observed in natural images Geisler et al. (2001).

### Neural dynamics

Inference in the presented model is realized by a biologically realistic dynamics in a network of neural populations that are linked by short- and long-range connections. This implementation of a dynamics is close to the approach of Rozell Rozell et al. (2008) but additionally includes long-range interactions between units with non-overlapping input fields. Most importantly, the constraint that only local visual information is available to the units receiving direct input from the visual field implies, and predicts, that inference is performed by *two* separate neural populations with activities ***a*** and ***b*** and different connection structures.

It is worth to speculate about a direct relation to the particular properties of neurons and anatomical structures found in different layers and between areas of visual cortex: Physiological studies distinguish between the near (*<* 2.5 degrees) and far surround (*>* 2.5 degrees) in contextual modulation Angelucci et al. (2017). Taking into account the spread of long-range horizontal axons within V1, which is less than about three degrees in visual space Angelucci et al. (2002), it seems likely that near surround effects are predominantly caused by horizontal interactions, while far surround effects are rather explained by feedback from higher visual areas. Assuming that one input patch in the model spans across 3 degrees in visual space, which is not implausible given the spatial extent of Gabor-like input fields shown in Fig. 3**A** (up to 1 degree in cortex), we would therefore identify ‘local’ interactions *W* ^local^ = −Φ^T^Φ with horizontal axons within V1, while ‘long-range’ interactions *W* ^long^ = *C* would be mediated by the combination of feedforward and feedback connections between visual cortical areas. A possible circuit diagram emerging from this paradigm is depicted in the left scheme in Fig. 8.

**Figure 8.**
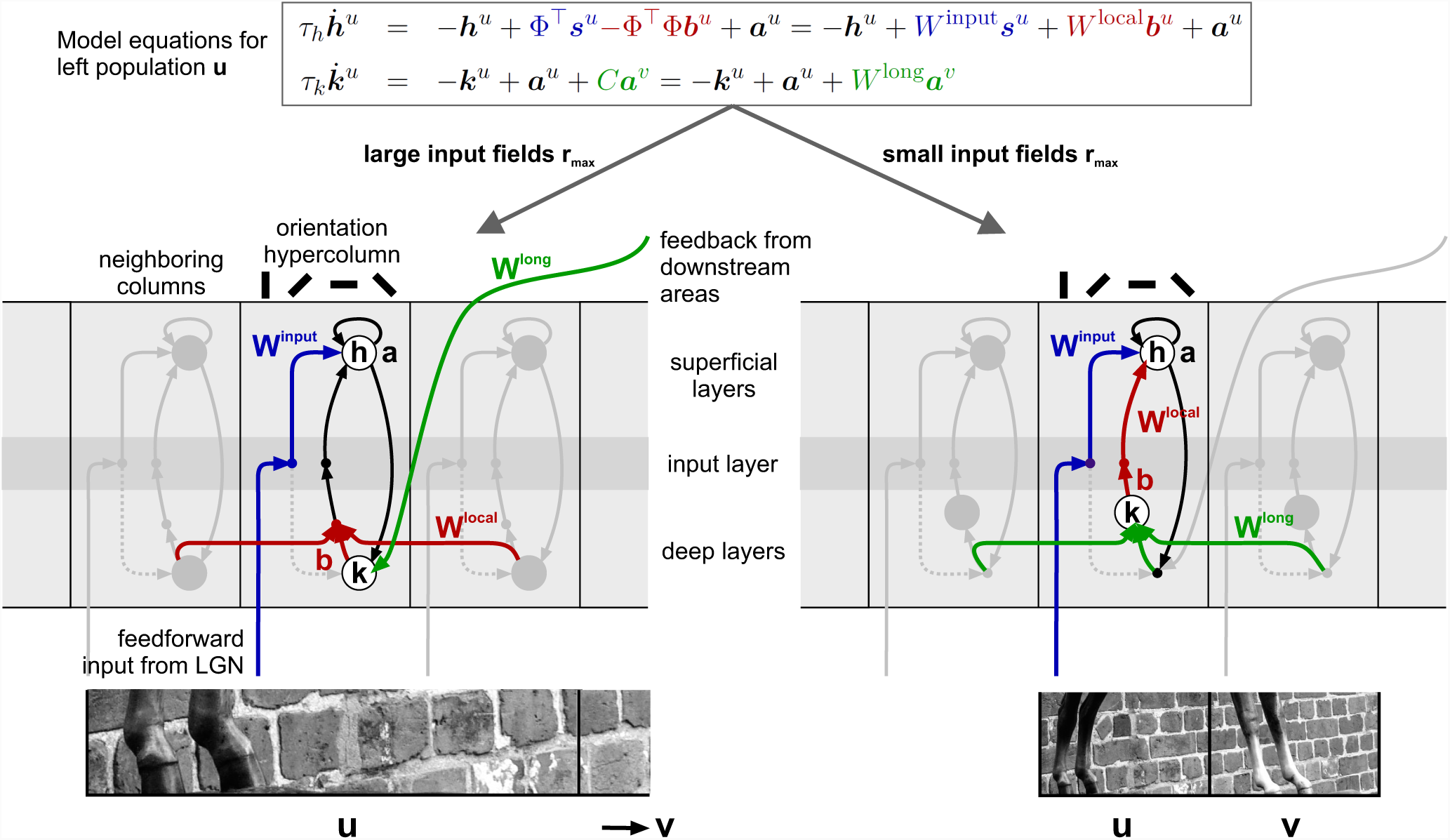
Putative neural circuits performing inference in visual cortex. Depending on the assumed spatial scale of input fields in the generative model, one distinguishes between cortical circuits where ‘long-range’ interactions *W* ^long^ would be mediated by recurrent loops between different cortical layers and ‘local’ interactions *W* ^local^ by long-ranging horizontal axons within primary visual cortex **(left scheme)**, or where long-range interactions *W* ^long^ would be mediated by long-ranging horizontal axons, and local interactions *W* ^local^ by the dense vertical/horizontal connection structures within a cortical hypercolumn **(right scheme)**. The length scales of input fields are indicated by the size of the image patch sections shown below. Interaction pathways associated with *W* ^long^, *W* ^local^ and *W* ^input^ are indicated in green, red and blue, respectively. Other links realizing different parts of the model equations (above the schemes) for column *u* are drawn in black. The putative connection schemes are embedded into sections of primary visual cortex with light and dark gray shading indicating different layers. Note that in the real cortex, also other connections such as links between input layer IV and deep layers (dashed, in gray) exist, and that interactions might be indirect by being relayed over intermediary target populations (filled dots) such as inhibitory interneurons.

An alternative picture evolves if we assume that input patches correspond to smaller regions in visual space. Now horizontal interactions within V1 would span over sufficiently long distances to mediate long-range interactions in the model (*W* ^long^ = *C*), while local interactions *W* ^local^ would indeed be local to a cortical (hyper-)column, possibly realized by the dense network linking different cortical layers in a vertical direction (example circuit shown in Fig. 8, right scheme).

In both discussed scenarios structure and polarity of cortical interactions is compatible with the model: horizontal and feedback connections are orientation-specific, and their effective interaction can be positive or negative Hirsch and Gilbert (1991); Weliky et al. (1995) since they have been found to target both, excitatory and inhibitory neurons McGuire et al. (1991).

### Contextual effects

Consistently the model reproduces a large variety of contextual phenomena, including size tuning, orientation-contrast effects and luminance-contrast modulations. In particular, all classical and non-classical receptive fields emerge in a fully unsupervised manner by training the model with ensembles of natural images. After training is finished, reproduction of all reported results is possible without change or fine-tuning of parameters, gains or thresholds – just by adhering to the exact visual stimulation procedures as used in the corresponding experimental studies. It is intriguing that also variability of the observed phenomena is reliably reflected in the statistics of model responses. This close match to experimental findings indicates that the assumed constraints from which dynamics and structure of the model were derived are constructive for providing a comprehensive framework for contextual processing in the visual system.

The nature of the observed effects, being orientation-specific and exhibiting both enhancement and suppression (see Figs. 5, 6, 7), closely mirrors the structures and polarities of local and long-range interactions. Furthermore, they explicitly link functional requirements to the anatomy of the visual system: As already observed in Zhu and Rozell (2013), local interactions between similar features are strongly suppressive. They realize competition between alternative explanations of a visual scene which is related to ‘explaining away’ in Bayesian inference Lochmann et al. (2012). The effects of long-range interactions depend on the exact stimulus configuration, and on the balance between neural thresholds and the combination of all recurrent inputs in the inference circuit. They serve to integrate features across distances, leading to the enhancement of noisy evidence such as in low-contrast stimuli Polat et al. (1998), but also to the suppression of activation by the model finding a simpler explanation for a complex stimulus configuration (i.e., by expressing the presence of multiple collinear line segments in terms of a single contour). This explicit link of natural statistics and cortical dynamics to function is also reflected in psychophysical studies: For example, in natural images an edge co-occurrence statistics being similar to correlation matrix *C* was observed and used to quantitatively predict contour detection performance by human subjects via a local grouping rule Geisler et al. (2001). High-contrast flankers aligned to a low-contrast center stimulus strongly modulated human detection thresholds Polat and Sagi (1993), providing facilitation over long spatial and temporal scales of up to 16 seconds Tanaka and Sagi (1998). Also detection thresholds of 4-patch stimulus configurations are closely related to natural image statistics Ernst et al. (2016). In both Ernst et al. (2016); Polat and Sagi (1993), the interactions between feature detectors with similar cRF properties are inhibitory for near contexts, and exhibit disinhibitory or even facilitatory effects for far contexts – paralleling the differential effects that local and long-range interactions have in our model.

In parallel to sparse coding, hierarchical predictive coding has emerged as an alternative explanation for contextual phenomena Rao and Ballard (1999). The general idea is that every layer in a cortical circuit generates an error signal between a feedback prediction and feedforward inference, which is then propagated downstream in the cortical hierarchy. While being conceptually different on the inference dynamics, the corresponding hierarchical generative model of visual scenes is similar to our paradigm when subjected to spatial constraints.

Besides principled approaches, contextual processing has been investigated with models constructed directly from available physiological and anatomical evidence Stemmler et al. (1995); Somers et al. (1998); Schwabe et al. (2006). Core circuit of such models is often an excitatory-inhibitory loop with localized excitation and broader inhibition and different thresholds for the excitatory and inhibitory populations, which is similar to our proposed cortical circuits shown in Fig. 8 with self-excitation of ***a*** and direct excitation on ***b*** and broader inhibition provided by *W*^local^ back onto ***a***. Such local circuits are connected by orientation-specific long-range connections, similar to the connections represented by *W* ^long^, even though they are typically assumed to be more strongly tuned. From these structural similarities we would speculate that contextual effects are caused in both model approaches by similar effective mechanisms.

### Outlook

In summary, our paradigm provides a coherent, functional explanation of contextual effects and cortical connection structures from a first-principle perspective, which requires no fine-tuning to achieve a qualitative and quantitative match to a range of experimental findings. For future studies, the model has some important implications:

First, there are experimentally testable predictions. These include the strong dependency of local and long-range interactions on the relative phase of adjacent classical receptive fields. Furthermore, we find two structures emerging in matrix *C*, namely a diagonal indicating stronger links between neurons with similar orientation preferences, as known from the literature, but also an anti-diagonal indicating enhanced links between neurons with opposite orientation preferences. Since connection probabilities were always reported w.r.t. orientation differences, the latter effect awaits experimental validation. Finally, we expect differences in the statistics of contextual effects between representations ***a*** and ***b*** to show up when information about the laminar origin of neural recordings is taken into account.

Second, it is formally straightforward to go back from the simplified model with just two separate input fields to the spatially extended, general scheme and subject it to much ‘broader’ visual scenes. Moreover, the neural dynamics allows also to address temporal contextual effects, or how neurons would respond to temporally changing contexts in the stimulus such as in ‘natural’ movies. For example, in simulations we observed strong transient effects shortly after stimulus onset (not shown), but a more thorough investigation and comparison to physiological findings is beyond the scope of this paper.

## Methods

### Learning and analysis of Φ and *C*

Variables Φ and *C* were learned using the procedure outlined in the Results section (eqs. (9)-(10)). We sampled input patches of size 16 *×* 32 pixels from a database of natural images Olmos and Kingdom (2004) from which we selected 672 images of size 576 *×* 768 pixels in uncompressed TIFF format. Images were first converted from RGB color space to grayscale values and then whitened using a method described in Olshausen and Field (1997). The optimization step for ***a*** (eq. (9)) was carried out for a batch of 100 image patches with a learning rate of *η*_*a*_ = 0.01. At the end of each update step for Φ (eq. (10)), the columns of Φ were normalized such that ||Φ_*i*_||_2_ = 1. We learned *N* = 1024 feature vectors. Learning was performed with 10^4^ iterations each for Φ and *C* (choosing as learning rates the values *η*_Φ_ = 0.05 and *η*_*C*_ = 0.01), after which both dictionary and long-range correlation matrix were stable. To obtain a better statistics, we repeated learning of the dictionary and of the long-range interactions several times, initializing the simulations with different seeds. The results presented in Figs. 4-7 are based on *N*_seed_ = 8 instances of the model.

To parametrize the feature vectors in terms of orientation, spatial frequency, size and location we fitted to each of them a Gabor function of the form

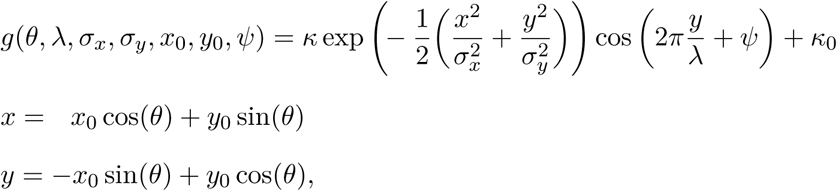

where *θ* is the orientation of the sinusoidal carrier, *λ* its wavelength, *ψ* its phase, *σ*_*x*_ and *σ*_*y*_ are the standard deviations of the gaussian envelope, *κ >* 0 the contrast and *κ*_0_ an offset. Fitting was done following a standard least square approach.

### Simulation of the neural model

The four differential equations that define the neural model (eqs. (21)) were solved numerically with a Runge-Kutta method of order 4 for a time interval of *T* = 600 ms. The time constants *τ*_*h*_ and *τ*_*k*_ were chosen to be 10 ms, close to physiological values of neurons in cortex Dayan and Abbott (2001). For analyzing the responses, we discarded the initial transients and averaged over single cell activities over the last 333 ms, a period of time that allowed a complete cycle of the stimulus drifting with a temporal frequency of 3 Hz, being the average preferred speed for cortical neurons Foster et al. (1985).

To ensure positivity of neural responses, in addition to the differential equations eqs. (21) we had to introduce a linear threshold operation (eqs. (12) and (18)). In contrast, no constraint is imposed on the sign of ***a*** and ***b*** in the generative model (eqs. (4)-(7)), nor by the optimization equation (9). To make the neural model consistent with the generative model, we therefore duplicated the number of neurons by introducing ON- and OFF units (see subsection Inference with a biologically plausible dynamics). In addition, we considered for all dictionary elements Φ_*i*_ also their mirrored versions −Φ_*i*_ and we split the long-range interactions into positive and negative contributions *C*^+^ = max(*C*, 0) and *C*^*-*^ = min(*C*, 0) via

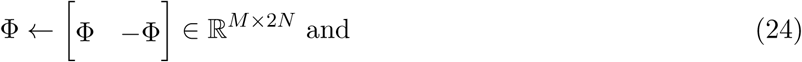

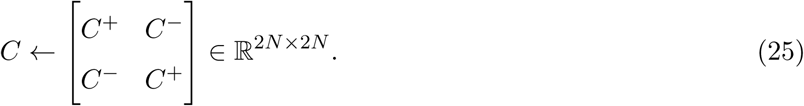

For selecting cells well-tuned and well-responding to stimuli centered in one input patch (see Contextual effects), all units were first stimulated with a set of small drifting sinusoidal gratings centered at ***r***^*u*^ with *r*_*c*_ = 2 pixels and *k*_*c*_ = 1. We varied *θ*_*c*_ from 0 to *π* in steps of *π/N*_*θ*_ (*N*_*θ*_ = 36) and the spatial frequency *f*_*c*_ from 0.05 to 0.35 cycles/pixel in steps of 0.025. We then selected for each neuron the preferred orientation and preferred spatial frequency. A unit was said to be responsive if its peak response was at least 10% of the maximum recorded activity. We determined orientation selectivity by computing, for each unit *n*, the complex vector average

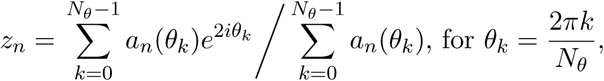

and we considered tuned those neurons for which it was |*z*_*n*_| *>* 0.85, corresponding to a tuning width of approximately 20 degrees half-width. With these selection criteria, we were left with 490 cells from all *N*_seed_ instantiations of the model.

### Selection of orientation contrast tuning classes

When we quantified the effect of cross-orientation stimulation, we pooled responses of units exihibiting the same qualitative behavior (Fig. 6**C,D**). To determine which behavior a unit showed we first computed, for each unit *n* with preferred orientation *θ\**, the average response to the compound stimulus when the surround orientation was close to *θ\**

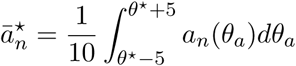

and when the surround orientation was near-oblique

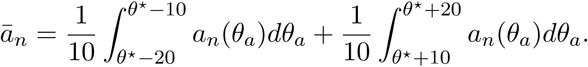

The unit was considered to show iso-orientation suppression if 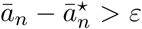, release from suppression if 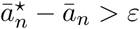 and untuned suppression in all other cases (*ε* = 0.05).

### Constants and parameters

Parameters used in numerical simulations are summarized in Table 1.

**Table 1.** Parameter values.

| Size tuning |  |  |  |
| --- | --- | --- | --- |
| $\theta_c$ | preferred | $\theta_a$ | - |
| $\omega_c$ | preferred | $\omega_a$ | - |
| $r_c$ | from 2 to 32 in steps of 1 | $r_a$ | - |
| $k_c$ | 1 | $k_a$ | - |
| Orientation-contrast (center-only) |  |  |  |
| $\theta_c$ | from 0 to $\pi$ in steps of $\pi/36$ | $\theta_a$ | - |
| $\omega_c$ | preferred | $\omega_a$ | - |
| $r_c$ | optimal | $r_a$ | - |
| $k_c$ | 1 | $k_a$ | - |
| Orientation-contrast (center-surround) |  |  |  |
| $\theta_c$ | preferred | $\theta_a$ | from 0 to $\pi$ in steps of $\pi/36$ |
| $\omega_c$ | preferred | $\omega_a$ | preferred |
| $r_c$ | optimal | $r_a$ | ∞ |
| $k_c$ | 1 | $k_a$ | 1 |
| Luminance-contrast (center-only) |  |  |  |
| $\theta_c$ | preferred | $\theta_a$ | - |
| $\omega_c$ | preferred | $\omega_a$ | - |
| $r_c$ | optimal | $r_a$ | - |
| $k_c$ | from 0.1 to 1 in steps of 0.1 | $k_a$ | - |
| Luminance-contrast (center-surround) |  |  |  |
| $\theta_c$ | preferred | $\theta_a$ | preferred |
| $\omega_c$ | preferred | $\omega_a$ | preferred |
| $r_c$ | optimal | $r_a$ | ∞ |
| $k_c$ | from 0.1 to 1 in steps of 0.1 | $k_a$ | 1 |

## Acknowledgments

We would like to thank Andreas K. Kreiter for fruitful discussions about the potential links of our model framework to physiology and anatomy of the primate visual system.

